# The causal role of early visual cortex in vividness of visual imagery

**DOI:** 10.1101/2024.04.25.591061

**Authors:** Juha Silvanto, Silvia Bona, Zaira Cattaneo

## Abstract

While numerous studies have demonstrated objective accuracy in visual imagery tasks to involve the early visual cortex (V1/V2), the role of this region in imagery vividness remains unclear. We addressed this question by combining Transcranial Magnetic Stimulation (TMS) with a cross-adaptation paradigm in which visual adaptation modulates the ability to engage in visual imagery in the adapted part of the visual field. As previously shown, in the control (Vertex) condition, mental imagery accuracy was impaired when the mental image was generated in the adapted region of visual space. This effect was removed by TMS, indicating that the locus of adaptation was the early visual cortex. In contrast, analysis of vividness ratings (collected on trial-by-trial basis) showed no effects of adaptation, indicating that the imagery vividness is somewhat orthogonal to the fidelity of the mental image. The key finding was that TMS impacted vividness ratings on trials when participants performed incorrectly in the imagery task. Thus, the effects of TMS on accuracy and vividness were observed on different trial types: TMS impaired accuracy of trials associated with high baseline performance, but increased vividness on incorrect trials. Overall, our findings suggest that the early visual cortex plays a role in both accuracy and vividness of mental images, yet these processes are dissociable.

## Introduction

We use mental imagery in many activities in everyday life. For example, when we are going from one place to another, we can imagine our route. Or when looking for lost keys, we can imagine where we saw them last. Already Sir Francis Galton (1880) reported large individual differences even within neurologically healthy individuals in these abilities. On one end of the spectrum are people (appr. 5 % of the population) who experience mental images that are so vivid that they can confuse them with real images; on the other end, 2-3% of individuals report the inability to create mental images, a condition referred to as aphantasia (see Zeman, 2024).

Interestingly, individuals unable to experience mental images can nevertheless perform within the normal range in various neuropsychological tasks which are thought to rely on imagery (e.g. Keogh et al, 2018; Milton, 2021; Pounder et al, 2022; see Zeman, 2024 for review) and even in tasks explicitly requiring visual imagery (Jacobs et al, 2018). This raises the question whether the subjective experience of imagery (i.e. its vividness) and its underlying representation are distinct. For example, it has been proposed that conditions like aphantasia reflect distinct conscious and unconscious representations of visual imagery. If the latter is intact, imagery processes can be carried out without conscious experience of that image (Nanay, 2021). In similar vain, Jacobs and Silvanto (2015) proposed that the phenomenal experience of a mental image is based on a distinct representation from the underlying memory trace.

From neural perspective, neuroimaging (see e.g. Pearson, 2015 for review) and brain stimulation evidence (e.g. Cattaneo et al., 2009a; Kosslyn et al., 1999) has implicated a range of visual cortical areas with visual imagery, including the early visual cortex. With respect to imagery vividness, there have been only few studies and the results are conflicting. Bergmann et al (2016) found a positive relationship between V1 volume and imagery precision (spatial location and orientation) whereas subjective vividness of imagery was positively related to prefrontal cortex volume. In contrast, Tabi et al (2022) found a small but significant positive correlation between the V1 volume and VVIQ scores. In terms of brain activity, Koenig-Robert and Pearson (2019) found that activity patterns in the primary visual cortex (V1) measured before a stimulus-related decision was performed, predicted future imagery vividness.

Here we used TMS to investigate the causal role of the early visual cortex (V1/V2) in imagery vividness. This was done by combining TMS with a visual adaptation paradigm. Behaviourally, adaptation manifests as impaired performance when the adapter and test stimuli share the same neuronal representation. Following visual adaptation, online TMS can remove its effect if the targeted region contains neuronal representations essential for encoding that feature (see e.g. Hartwigsen & Silvanto, 2022; Silvanto & Cattaneo, 2017; Silvanto et al., 2018). In this manner, in combination with adaptation, TMS can be used to assess neuronal representations in specific brain areas. In our previous study (Cattaneo & Silvanto, 2012), mental imagery accuracy was found to decrease when the spatial location for generating the mental image had been adapted through visual stimulation. The application of TMS over the early visual cortex abolished the effects of visual adaptation, indicating that this region is a locus of the adaptation effect.

To examine the neural basis of imagery judgments, we asked participants provide a vividness rating of their mental image on trial-by-trial basis, in addition to performing the discrimination task. As in the previous study, TMS was applied over the early visual cortex, in order to determine whether vividness ratings are similarly affected by adaptation and TMS as accuracy ratings.

## Material and Methods

### Participants

21 right-handed participants (7 males, mean age = 23.5 years, SD = 2.28) with normal or corrected-to-normal visual acuity were recruited for the study. Prior to their inclusion in the study, each participant was screened for contraindication for TMS and provided a written informed consent. The study was approved by local ethics committee and all participants were treated in accordance with the Declaration of Helsinki.

### Experimental procedure

#### Stimuli and task

The paradigm was adapted from the study of Cattaneo & Silvanto (2012). Stimuli were presented using E-Prime Software 2.0 (Psychology Software Tools Inc., Pittsburgh, USA) centrally on a 15.5-inch monitor (1200× 800 resolution with a viewing distance of 60 cm). Fig 1 shows an example of an experimental trial. At the start of each trial, a top-up adapter lasting for 5 seconds was presented (please see next section for details of adaptation conditions). This was followed by fixation cross and the imagery cue. The latter consisted of a digital time in black digits (diameter of each digit being appr 1° of visual angle) and it was presented in the middle of the screen for 1 sec. A total of 36 digital times were used (18 corresponding to clock hands in the upper half of the clock face, and 18 in the bottom half). This was followed by a blank screen (200 ms) and black circle at fixation (with a diameter of 12° of visual angle; extending 6° into both hemifields). Participants task was to imagine the clock hands representing the digital time within the circle. At the end of the imagery phase, a target (a black dot with a diameter of 0.5° of visual angle) was presented in one of the four quadrants of the circle. Participants’ task was to judge whether the target appeared inside or outside the area defined by the hour and the minute hand of the clock (Cattaneo & Silvanto, 2012). In half of the trials it appeared inside this area and in the on the other half outside. On each trial, the target appeared 1° of visual angle away from the minute hand. In half of the trials the dot appeared in the upper part of the circle, and on the other half in the lower part of the circle; furthermore, in half of the trials the dot appeared on the left side of the circle, and on the other half on the right side of the circle. The position of the dot within the clock face appeared in random order and participants were instructed to maintain fixation at the center of the circle throughout the block. After their response, participants were asked to rate the subjective vividness (using a 1-4 scale) of the mental image of the clock hands. In this scale, “1” referred to the absence of a mental image and “4” referred to a mental image as clear and vivid as the visual perception of the clock hands. In each trial, 1000 ms after the onset of the imagery cue, participants received the TMS stimulation (see below).

**Figure 1.**
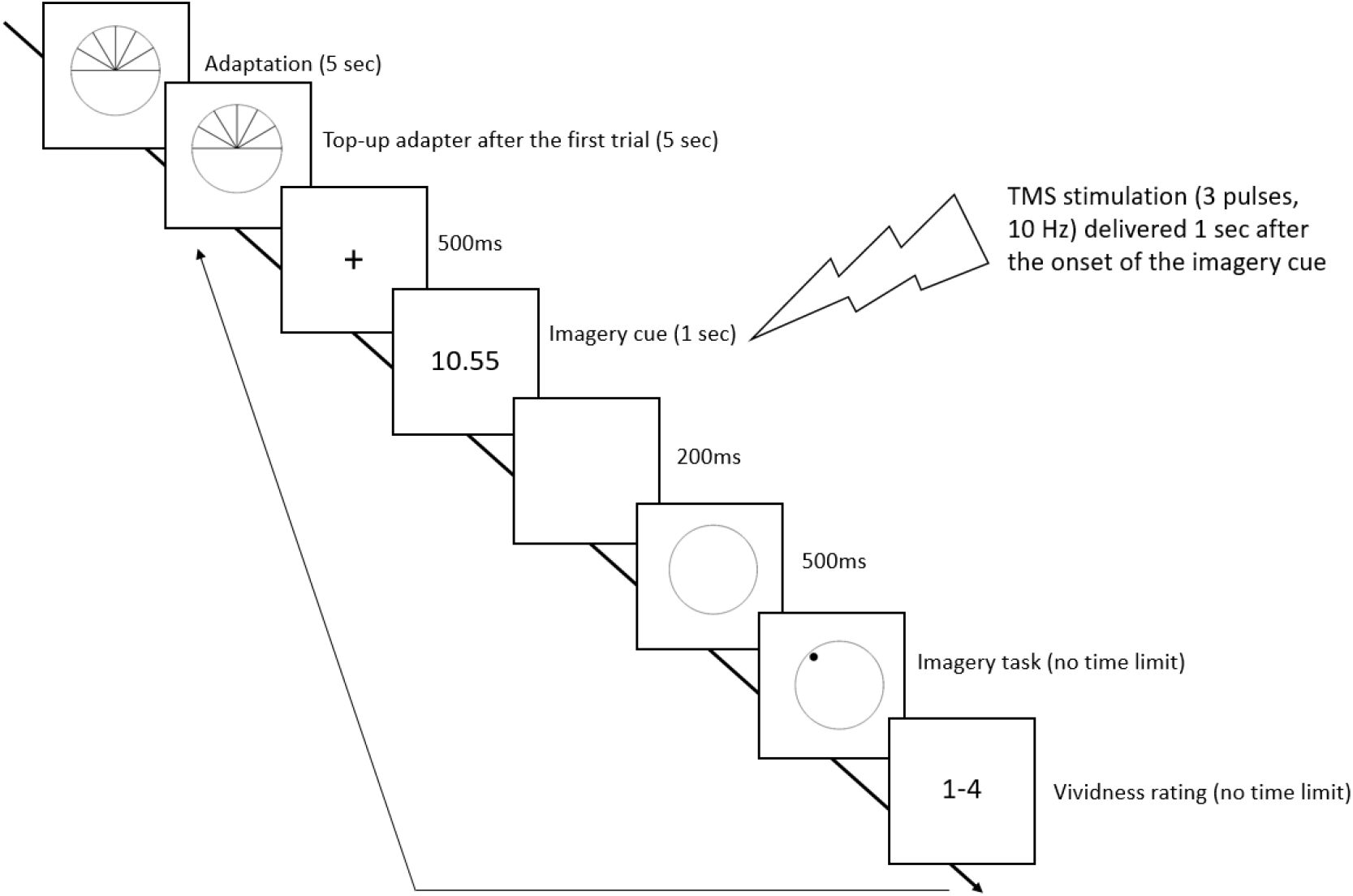
Timeline of an experimental trial. During each trial, participants were presented with an imagery cue, represented by a digital time, and instructed to mentally visualize the corresponding position of the clock hands within the clock face. After a delay of 500 milliseconds, a black dot appeared within the clock face, prompting participants to determine whether it fell inside or outside the area defined by the clock hands. Subsequently, participants were asked to rate the subjective vividness of their mental image using a 4-point scale. At the onset of each block, a 5-second adapter was displayed, and before each trial, a top-up adapter was presented for 5 seconds. Transcranial magnetic stimulation (TMS), consisting of 3 pulses at 10 Hz, was administered 1 second after the onset of the imagery cue.

#### Visual adaptation

Visual adaptation was conducted prior to the beginning of each block. During adaptation, participants were exposed to visual lines displayed either in the top or bottom half of a clock face (see Fig. 1). Each adaptation sequence comprised 7 lines positioned to overlap with potential locations of clock hands that participants were instructed to imagine, situated either in the upper or lower half of the clock face. Within each experimental block, two adaptation loops were randomly presented: one for adaptation to the upper half of the clock and another for the lower half. During each block, the adapter remained visible on the screen for 50 seconds, followed by 12 trials wherein different digital times were presented. Thus, each block encompassed 24 trials, with 12 trials dedicated to each type of adapter. Among these trials, 6 involved mental imagery in the upper half of the clock, while the remaining 6 focused on the bottom half. Preceding all trials was a top-up adaptation phase, wherein the same adapter was displayed for 5 seconds. Each participant completed 6 blocks (2 blocks for each TMS condition, as described below), with each block containing 2 adaptation blocks (one for each adapter). Before the commencement of the experimental sessions, participants underwent a brief practice session (16 trials, with 8 trials for each type of adapter) to familiarize themselves with the task.

### TMS

TMS was applied with a Magstim Rapid2 stimulator (Magstim Co Ltd., Whitland, UK) using a 70-mm coil. The stimulation protocol consisted of 3 pulses applied at a frequency of 10 Hz (*i*.*e*., 100ms gap between the pulses). The intensity was fixed at 60% of stimulator output, an intensity often used in TMS studies on visual cortex (e.g. Campana et al, 2002). Stimulation was applied over the early visual cortex (V1/V2) and over the Vertex, 1000 ms after the onset of the imagery cue, aiming to interfere during the interval in which the digital time has been processed and the mental image is being created. V1/V2 was localized by placing the coil 2 cm above the inion, which is likely to correspond to the location of the early visual cortex (e.g. Thielscher et al, 2011). The Vertex was used as the baseline condition to control for non-specific, somatosensory effects induced by the stimulation. None of the participants reported perception of phosphenes during the experiment.

## Results

One participant withdrew from the experiment due to TMS-induced discomfort during stimulation and two further participants were excluded due to below-chance performance (< 50%) in the imagery task; therefore, the final sample included 18 participants (7 males, mean age = 23.53 years, SD = 2.46). Statistical analyses were performed separately for the imagery task and for the vividness rating. Data were analysed as a function of the overlap in spatial positions of the adapter and the mental image (*i*.*e*. “Spatial Overlap”, “No Spatial Overlap”).

### Imagery task accuracy

The results for accuracy are shown in Figure 2, as a function of the spatial overlap between the location of the mental image and the adapter in each TMS condition. A repeated-measures ANOVA with TMS site (Vertex, V1) and Spatial Overlap (spatial overlap, no spatial overlap) as within-subjects factors on mean accuracy revealed a significant main effect of Spatial Overlap (*F*(1,17)=7.93, *p*=.012, 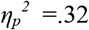) and a significant interaction TMS site by Spatial Overlap (*F*(1,17)=5.54, *p*=.031, 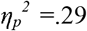). The main effect of TMS site was not significant (*F*(1,17)=2.51, *p*=.13, 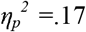). Post-hoc pairwise comparisons showed that stimulation of V1 impaired performance in no-spatial overlap trials compared to stimulation of Vertex (*t*(17)=2.24, *p*=.025) while having no impact on spatial overlap trials, *t*(17)=.413, *p*=.0.69). In summary, TMS was found to abolish the effects of adaptation.

**Figure 2.**
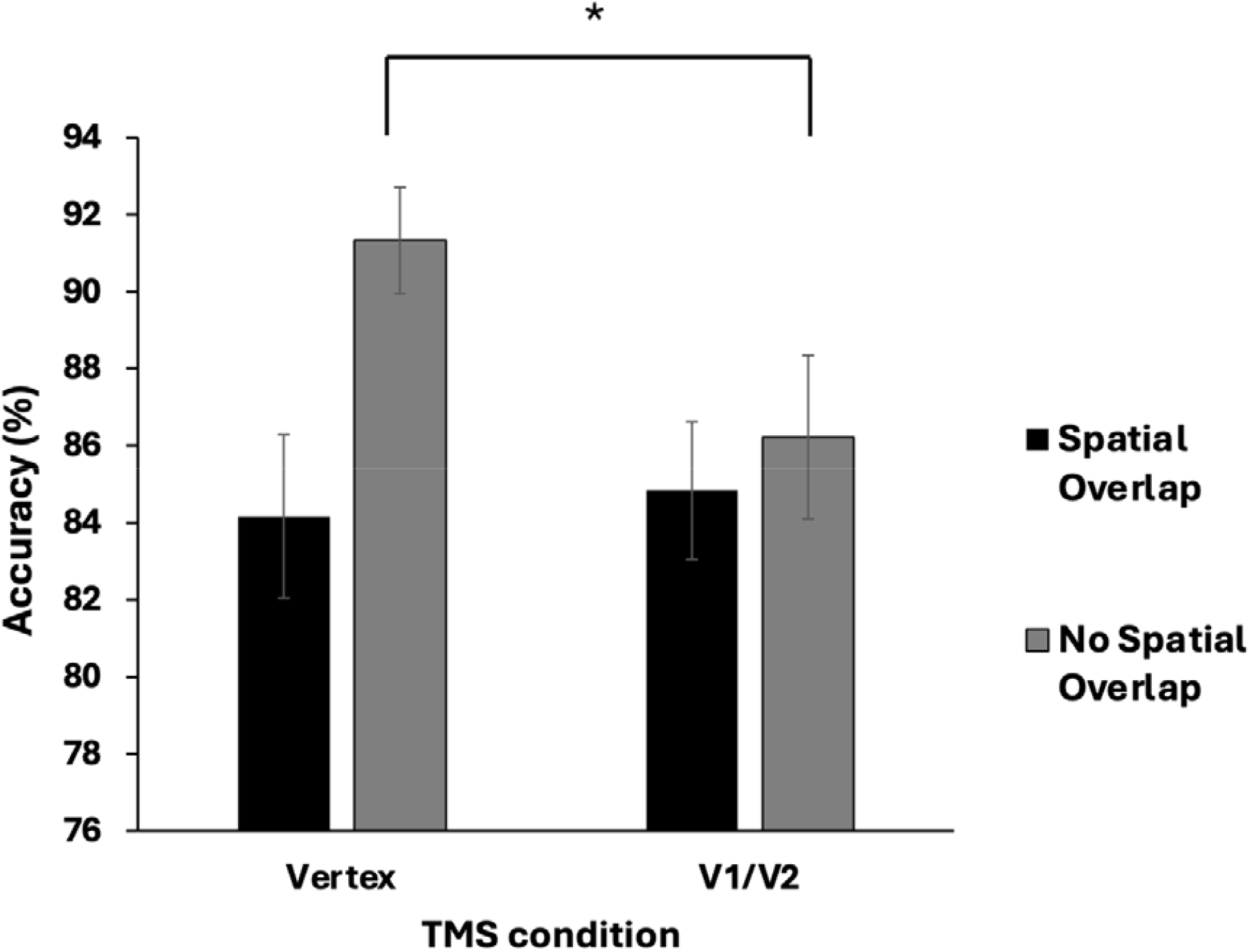
Mean (N=18) participants’ accuracy (%) as a function of TMS condition and Spatial Overlap between mental image and adapter. Compared to stimulation of Vertex, TMS over V1 disrupted detection of no-spatial overlap trials. In turn, perception of spatial overlap trials was unaffected. The asterisks indicate significant pairwise comparisons. Error bars reflect ±1 SEM.

### Vividness ratings

For vividness rating (shown in Figure 3), the same ANOVA found no significant effects of Spatial overlap (*F*(1,17)=0.46, *p*=0.51, 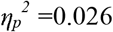) or TMS site (*F*(1,17)=0.003, *p*=.955, 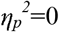) nor a significant TMS site by Spatial Overlap interaction (*F*(1,17)=.43, *p*=0.522, 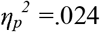). Thus it appears that there were no effects of either adaptation or TMS on vividness ratings.

**Figure 3.**
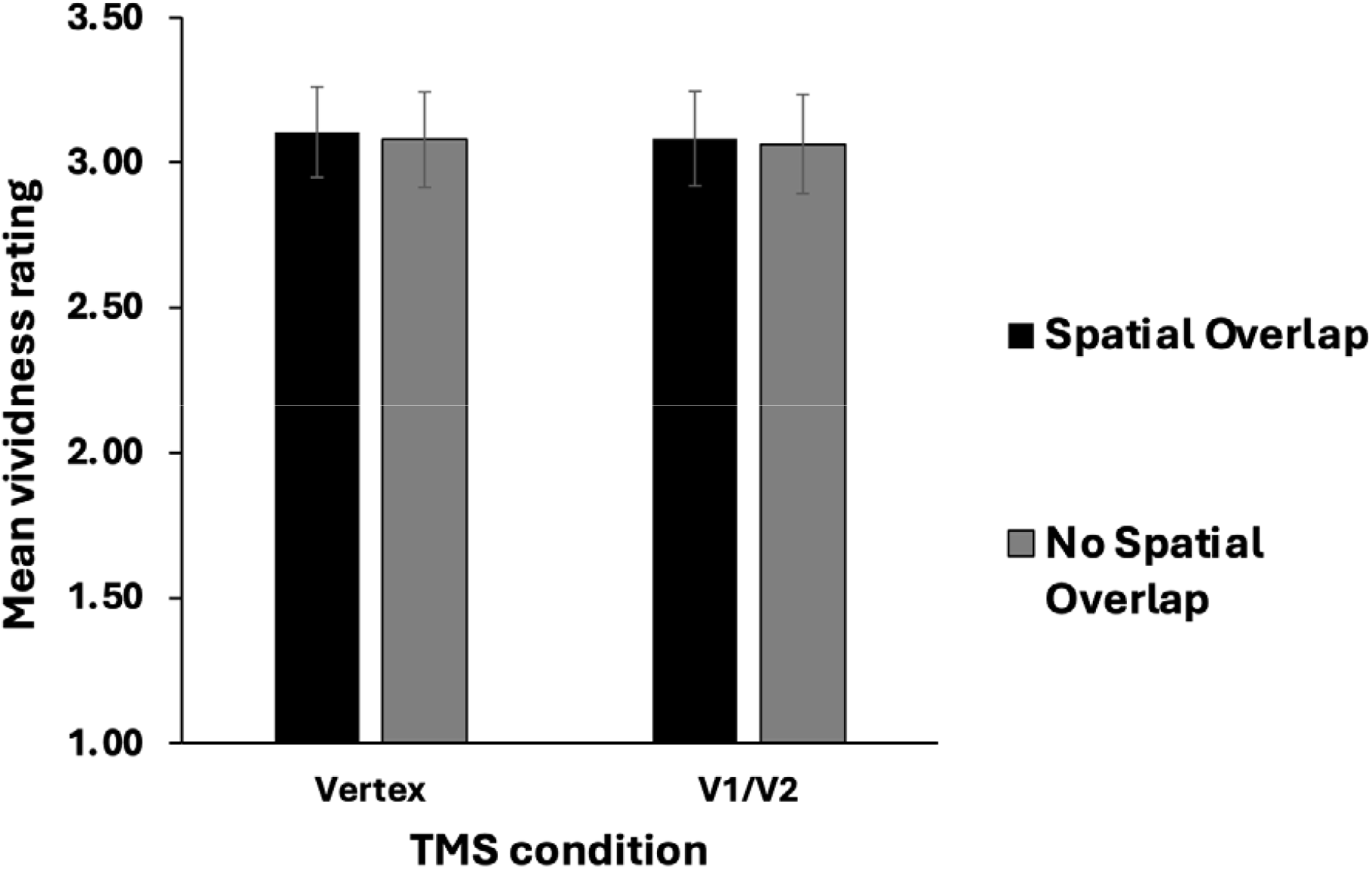
Mean vividness rating as a function of TMS condition and Spatial Overlap between mental image and adapter. Performance was not affected by either TMS or the Spatial Overlap. Error bars reflect ±1 SEM.

### TMS effects on accurate vs inaccurate trials

We then carried out an analysis where vividness ratings were analyzed as a function of whether they are associated with a correct or incorrect answer in the imagery task. The logic here is that TMS might have a differential effect depending on the vividness of the mental image, and less vivid mental images are generally associated with incorrect responses. These results are shown in Figure 4. We included an additional main factor into the ANOVA, namely response accuracy. ANOVA found no main effect of Spatial overlap (F(1,17)=4.236, p=0.055, ηp2 =0.199), no main effect of TMS site (F(1,17)=2.934, p=0.105, ηp2=0.147), and a significant main effect of response accuracy (F(1,17)=27.41; p<0.001; ηp2 =0.717). In addition, there was no significant interaction between Spatial overlap and TMS site (F(1,17)=0423; p=0.524, ηp2 =0.-24) or Spatial overlap and response accuracy (F(1,17)=3.24; p=0.09, ηp2 =0.160). In contrast, the interaction between TMS site and Response accuracy (F(1,17)=5.030 p=0.039, ηp2 =0.228) was significant. The three-way interaction between Spatial overlap, TMS site and Response accuracy was also statistically non-significant ((F(1,17)=0.370; p=0.551, ηp2 =0.021). In summary, the ANOVA shows that TMS facilitated vividness ratings for incorrect trials, independently of spatial overlap between visual adaptation and the mental image.

**Figure 4.**
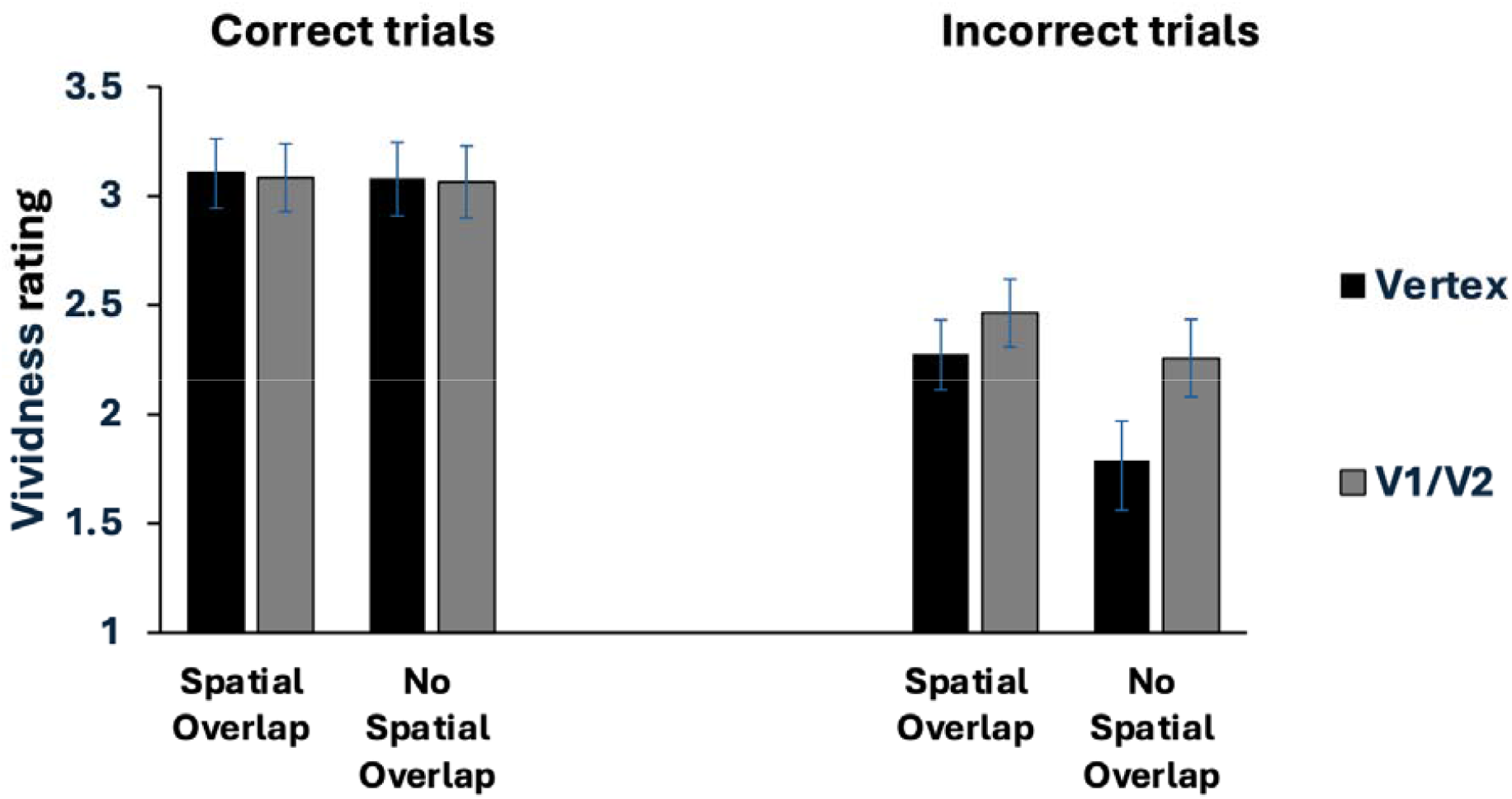
Mean vividness rating separately for correct and incorrect trials, as a function of TMS condition and Spatial Overlap between mental image and adapter. Performance was not affected by either the TMS effect or the Spatial Overlap. Error bars reflect ±1 SEM.

## Discussion

For imagery accuracy, performance was significantly lower when the mental image spatially overlapped with the adapter (i.e. “Spatial Overlap” condition) relative to when the mental image needed to be created in the other half of the clock face (i.e. “No Spatial Overlap” condition). Thus the adaptation paradigm induced the effect observed previously (Cattaneo & Silvanto, 2012). The application of TMS over the early visual cortex interacted with adaptation, such that the adaptation effect present in the baseline (Vertex) condition was abolished. Specifically, TMS impaired performance on “No Spatial Overlap” trials whereas “Spatial Overlap” trials were unaffected.

The main objective of this study was to examine the effects of TMS on vividness ratings. When these ratings were analyzed as a function of congruency to adapting stimulus, no impact of adaptation or of TMS were found (see Figure 3). This would lead one to conclude that vividness ratings are fully orthogonal to the accuracy mental image which underlies performance in this task. Furthermore, the conclusion would be that the imagery vividness, unlike the accuracy of that mental image, does not rely on early visual cortex.

However, the picture is more complicated when vividness ratings are analyzed as a function of accuracy (as shown in Figure 4). As one would expect, vividness ratings are lower for incorrect vs correct trials. The key finding is that TMS was found to increase vividness on incorrect trials (which were associated with a relatively lower baseline vividness) but not on correct trials (which had a higher baseline vividness). This facilitation was not dependent on adaptation condition (i.e. Spatial overlap).

This raises the question regarding the link between mental image accuracy and vividness. On one hand, one would expect the two to go together, as presumably more accurate mental representations (as assessed by task accuracy) would be expected to be more vivid. The pattern of results in the baseline (Vertex) condition is consistent with this. Interestingly, however, the TMS effect observed here indicates a dissociation between the two because TMS effects on accuracy and vividness involve different trials. Specifically, the effect of TMS on mental image accuracy was observed on “No Spatial overlap” trials; TMS did not modulate mental image accuracy on Spatial Overlap trials. In contrast, the effect of TMS on vividness was independent of Spatial Overlap. Rather, TMS enhanced vividness responses generally on incorrect trials. In short, TMS decreased accuracy when baseline accuracy was high, but increased vividness for inaccurate responses.

One might argue that the vividness ratings here would be better described as metacognitive ratings of task accuracy, rather than judgments of imagery vividness. In other words, perhaps participants, when realizing having made an error, use this as a heuristic for the subsequent vividness judgment, rather than judging vividness on its own merits. This issue can be somewhat difficult to disentangle as the two possibilities would not necessarily lead to a different behavioral pattern, given that generally a more accurate mental image would also likely be more vivid. However, prior studies provide evidence against this possibility. Bona et al (2013) showed that task accuracy and vividness ratings are differentially affected by distracters presented during the trial, indicating that the two judgments can be based on different sources of information. Our TMS results are also inconsistent with this view because accuracy and vividness were affected on different types of trial. Specifically, as TMS increased errors for the “Spatial Overlap” condition, this should have been associated with a reduction of vividness ratings in the same condition. However, this pattern of results was not observed. It is thus more parsimonious to assume that participants indeed followed the task instruction of judging imagery vividness.

The effects observed here are consistent with the literature on state-dependent TMS effects which has shown the direction of TMS effects to depend on the baseline performance level. There is much evidence to show that TMS-induced disruptions occur at relatively high baseline levels, whereas with lower levels of performance, facilitations can occur (see e.g. Hartwigsen & Silvanto, 2022; Silvanto & Cattaneo, 2017 for review). This is indeed what was found here, with TMS impairing accuracy when baseline accuracy level was high, and facilitating vividness rating when baseline vividness was low. The key finding is that these were not the same trial types – TMS effects on accuracy and vividness did not go hand in hand. Thus, while both the fidelity and vividness appear to causally involve early visual cortex, they appear to rely on at least partially distinct mechanisms.

There is very little prior evidence on the neural basis of imagery vividness. Perhaps the strongest evidence so far is that activity patterns in V1 could be used to decode imagery vividness many seconds before the actual imagery judgment (Koenig-Robert & Pearson, 2019). However, with respect to visually induced phenomenal awareness, there is a large literature, both empirical and theoretical, implicating V1 in the phenomenology of conscious experience (see e.g. Lamme, 2000; Dehaene & Naccache, 2001; Block, 2005, for reviews). This view has been supported also by more recent studies (e.g. Lyu et al, 2022). Our results would be consistent with the view that neural mechanisms that reflect conscious experience in the visual domain also contribute to the vividness of internally generated percepts.

The neural dissociation between the objective accuracy of mental images and their subjective vividness reported here is consistent with the nature of aphantasia. As mentioned in the Introduction, individuals who self-report having aphantasia show performance in cognitive tasks which does not differ from controls (see Zeman, 2024 for review). This dissociation suggests that the underlying imagery representation on which objective discrimination performance is based is intact in these individuals; the issue appears to be the inability to representation to conscious experience (Nanay, 2021). Our finding that accuracy and vividness were dissociable by TMS is consistent with this view.

In summary, our results provide causal evidence that the early visual cortex is involved in the subjective vividness of imagery, and that vividness and accuracy of the mental image rely on partly distinct mechanisms.

## References

Bergmann, J., Genç, E., Kohler, A., Singer, W., & Pearson, J. (2016). Smaller Primary Visual Cortex Is Associated with Stronger, but Less Precise Mental Imagery. Cerebral Cortex, 26(9), 3838–3850.

Bona S, Cattaneo Z, Vecchi T, Soto D, Silvanto J (2013). Metacognition of Visual Short-Term Memory: Dissociation between Objective and Subjective Components of VSTM. Front Psychol. 14;4:62. doi: 10.3389

Campana, G., Cowey, A., & Walsh, V. (2002). Priming of motion direction and area V5/MT: a test of perceptual memory. Cerebral Cortex, 12(6), 663–669.

Cattaneo, Z., Bona, S., & Silvanto, J. (2012). Cross-adaptation combined with TMS reveals a functional overlap between vision and imagery in the early visual cortex. Neuroimage, 59(3), 3015–3020.

Cattaneo, Z., & Silvanto, J. (2008). Time course of the state-dependent effect of transcranial magnetic stimulation in the TMS-adaptation paradigm. Neuroscience Letters, 443(2), 82–85.

Cattaneo, Z., Vecchi, T., Pascual-Leone, A., & Silvanto, J. (2009). Contrasting early visual cortical activation states causally involved in visual imagery and short-term memory. European Journal of Neuroscience, 30(7), 1393–1400.

Hartwigsen, G., & Silvanto, J. (2023). Noninvasive Brain Stimulation: Multiple Effects on Cognition. Neuroscientist, 29(5), 639–653.

Kosslyn, S. M., Pascual-Leone, A., Felician, O., Camposano, S., Keenan, J. P., Thompson, W. L., … & Alpert, N. M. (1999). The role of area 17 in visual imagery: convergent evidence from PET and rTMS. Science, 284(5411), 167–170.

Jacobs, C., & Silvanto, J. (2015). How is working memory content consciously experienced? The ‘conscious copy’ model of WM introspection. Neuroscience & Biobehavioral Reviews, 55, 510–519.

Dehaene S, and Naccache L. (2001). Towards a cognitive neuroscience of consciousness: basic evidence and a workspace framework. Cognition 79, 1–37.

Jacobs, C., Schwarzkopf, DS, Silvanto, J. (2018). Visual working memory performance in aphantasia. Cortex, 105, 61–73.

Keogh, R., Wicken, M., & Pearson, J. (2021). Visual working memory in aphantasia: Retained accuracy and capacity with a different strategy. Cortex, 143, 237–253.

Lamme, V. A., Supèr, H., Landman, R., Roelfsema, P. R., & Spekreijse, H. (2000). The role of primary visual cortex (V1) in visual awareness. Vision Research, 40(10-12), 1507–1521.

Milton, F., Fulford, J., Dance, C., Gaddum, J., Heuerman-Williamson, B., Jones, K., … & Huth, A. (2021). Behavioral and neural signatures of visual imagery vividness extremes: Aphantasia vs. Hyperphantasia. Cerebral Cortex Communications, 2(2), 1–15.

Nanay, B. (2021). Unconscious mental imagery. Philosophical Transactions of the Royal Society B: Biological Sciences, 376(1817), 20190689.

Pearson, J., Naselaris, T., Holmes, E. A., & Kosslyn, S. M. (2015). Mental Imagery: Functional Mechanisms and Clinical Applications. Trends in Cognitive Sciences, 19(10), 590–602.

Silvanto, J., Bona, S., Marelli, M., & Cattaneo, Z. (2018). On the mechanisms of Transcranial Magnetic Stimulation (TMS): How brain state and baseline performance level determine behavioral effects of TMS. Frontiers in Psychology, 9, 741.

Silvanto, J., & Cattaneo, Z. (2017). Common framework for “virtual lesion” and state-dependent TMS: The facilitatory/suppressive range model of online TMS effects on behavior. Brain and Cognition, 119, 32–38.

Tabi, Y. A., Maio, M. R., Attaallah, B., Dickson, S., Drew, D., Idris, M. I., … & Husain, M. (2022). Vividness of visual imagery questionnaire scores and their relationship to visual short-term memory performance. Cortex, 146, 186–199.

Thielscher, A., Reichenbach, A., Uğurbil, K., & Uludağ, K. (2010). The cortical site of visual suppression by transcranial magnetic stimulation. Cerebral Cortex, 20(2), 328–338.

Zeman, A. (2024). Aphantasia and hyperphantasia: exploring imagery vividness extremes. Trends in Cognitive Sciences. Advance online publication. doi: 10.1016/j.tics.2024.02.007.

